# Levonorgestrel modulates innate immune activation in human endocervical epithelium and monocytes during *Chlamydia trachomatis* infection

**DOI:** 10.1101/2019.12.16.878710

**Authors:** Alison J. Eastman, Sophia Liu, Jason D. Bell

## Abstract

**Problem:** Our group has previously shown that baboons with a levonorgestrel-releasing intrauterine system (LNG-IUS) have delayed clearance of *Chlamydia trachomatis* (Ct). Based on this result, we hypothesized that LNG results in changes to development of the immune response by epithelial and resident innate immune cells.

**Method of Study:** Using the end1 endocervical cell line or the THP.1 monocyte-like cell line, cells were exposed to increasing levels of progesterone (P4) or a dose of LNG representative of LNG in reproductive tract tissues of women with an LNG-IUS. Ct was used at an MOI of 1 and supernatants were collected for ELISA at 48 hours post-infection. Select nuclear receptors were inhibited to determine which receptor contributed to LNG-mediated immunosuppression.

**Results:** Cervical epithelial cells infected with Ct expressed IL-1β when treated with vehicle control. P4 further increased IL-1β expression during Ct infection, while LNG decreased IL-1β expression. Treatment with the androgen receptor blocker ailanthone prevented LNG-mediated immunosuppression.

**Conclusions:** LNG in the presence of increasing P4 suppresses IL-1β production in response to Ct infection *in vitro*. This appears to be mediated at least in part by the androgen receptor. This has implications for women with LNG-IUS at high risk for sexually transmitted infections.

## Introduction

Intrauterine devices (IUDs) are the most effective, long-acting, reversible contraception available to women worldwide^1–3^. For decades, many practitioners have limited the use of IUDs to women who have had children and are in monogamous, long-term relationships because of concerns that sexually transmitted infections (STIs), particularly *Chlamydia trachomatis* (Ct), may be more severe when an IUD is in place^4^. However, there has been a renewed interest in the use of the IUD in the US ^5^ and for use in populations that may be at a higher risk for unintended pregnancy and maternal mortality. The levonorgestrel-intrauterine system (LNG-IUS) was FDA approved in 2000^3,5–7^; however, the effects of LNG-IUS on STIs like Ct are largely unknown. Our group has shown that in a baboon model of Ct, the presence of LNG-IUS resulted in delayed clearance of Ct from the lower reproductive tract^8^. This may have consequences for women with the LNG-IUS. It is critically important to understand the immunologic or physiologic mechanisms behind LNG-IUS-associated delay in Ct clearance.

The global burden of Ct infection is enormous, with the World Health Organization reporting over 130 million cases each year^9^. The initial site of Ct infection is the cervical columnar epithelial cells, which begin to produce inflammatory cytokines in response to infection^10^. Interleukin-1β is produced in response to Ct infection by epithelial cells and phagocytes and is tightly regulated at many levels: epigenetically, transcriptionally, and post-translationally via activation of the inflammasome. The inflammasome is a multi-subunit protein complex that includes members of the nod-like receptor (NLR) family of cytosolic pattern recognition receptors (PRRs), which assemble with scaffold proteins to cleave pro-caspase-1 into active caspase-1, which then cleaves pro-IL-1β into active IL-1β^11^. Priming of NLRs occurs via the signaling pathways downstream of Toll-like receptors (TLRs) and other PRRs. Inhibition of the inflammasome pathway is known to occur by glucocorticoid signaling, which can be activated by promiscuous binding of progestins used in hormonal contraceptives. However, the inflammasome may act as a double-edged sword during Ct infection: while production of IL-1β is an early activator of the immune system and stimulates production of IL-8 (neutrophil chemoattractant)^12^ and CCL3 (T cell chemoattractant)^13,14^, it is associated with upper female reproductive tract pathology^15^, and activated inflammasome components can aid Ct in survival and proliferation in cytoplasmic inclusions^16^.

Progestogens- including progesterone (P4) and the synthetic progestogen (progestin) medroxyprogesterone acetate (MPA)- are immune modulators^17^. Distinct progestogens can act differently based on which nuclear receptors they stimulate and with what affinity (*e.g.* progesterone receptor (PR), glucocorticoid receptor (GR))^18–20^. P4 is known to be immunomodulatory, but this is both concentration and cell type-dependent; while frequently immunosuppressive, in high concentrations (≥0.1 μM) it can be pro-inflammatory^21^. P4 can facilitate IL-1β production and concomitant downstream inflammatory responses^22^. MPA is associated with lower expression of IL-1β during Ct infection^23^. Progestogens are known to affect TLR signaling^24^, and are hypothesized to alter inflammasome components^25,26^, although this has not been definitively shown. There is controversy surrounding LNG and whether it is immunomodulatory similar to MPA^27^. The LNG-IUS is known to alter immune cell populations in the female reproductive tract^27^, and high levels of systemic LNG have been shown to affect circulating immune cell numbers, which are further modulated by the duration of LNG contraception^28,29^.

To date, mechanistic studies of the immunomodulation of LNG have not considered how P4 may influence the effect of LNG on the immune system, particularly during Ct infection. In this study, we treated human endothelial cells and immune cells with increasing, biologically-relevant concentrations of P4 in the presence or absence of LNG and examined the impact on IL-1β production.

## Results and discussion

During use of an LNG-releasing intrauterine system (LNG-IUS), circulating levels of P4 range from ~0-1500 nM^30^. To model this, we subjected the endocervical epithelial cell line End1 ^31^ to increasing levels of P4 (0-1 μM) in the presence or absence of LNG at the concentration to which it accumulates in the uterus (808 ng/g tissue, 6937 pg/mg protein; extrapolated from Nilsson 1982^32^ to 161 ng/mL or 512 nM, which informed our use of LNG at 100 ng/mL). By qPCR on mRNA isolated from End1 cultures, Ct infection generally suppressed mRNA expression of the inflammasome component ASC during all concentrations of P4 treatment (Fig. 1A). However, this suppression was dramatically enhanced in the presence of LNG and P4 combined at all concentrations. P4 increased IL-1β mRNA in a dose-dependent manner during Ct infection, but addition of LNG suppressed IL-1β mRNA levels to baseline (Fig. 1B). IL-1β protein was induced by P4 during Ct infection, but the addition of LNG suppressed that at high levels of P4 treatment (100 and 1000 nm P4) (Fig. 1C).

**Figure 1.**
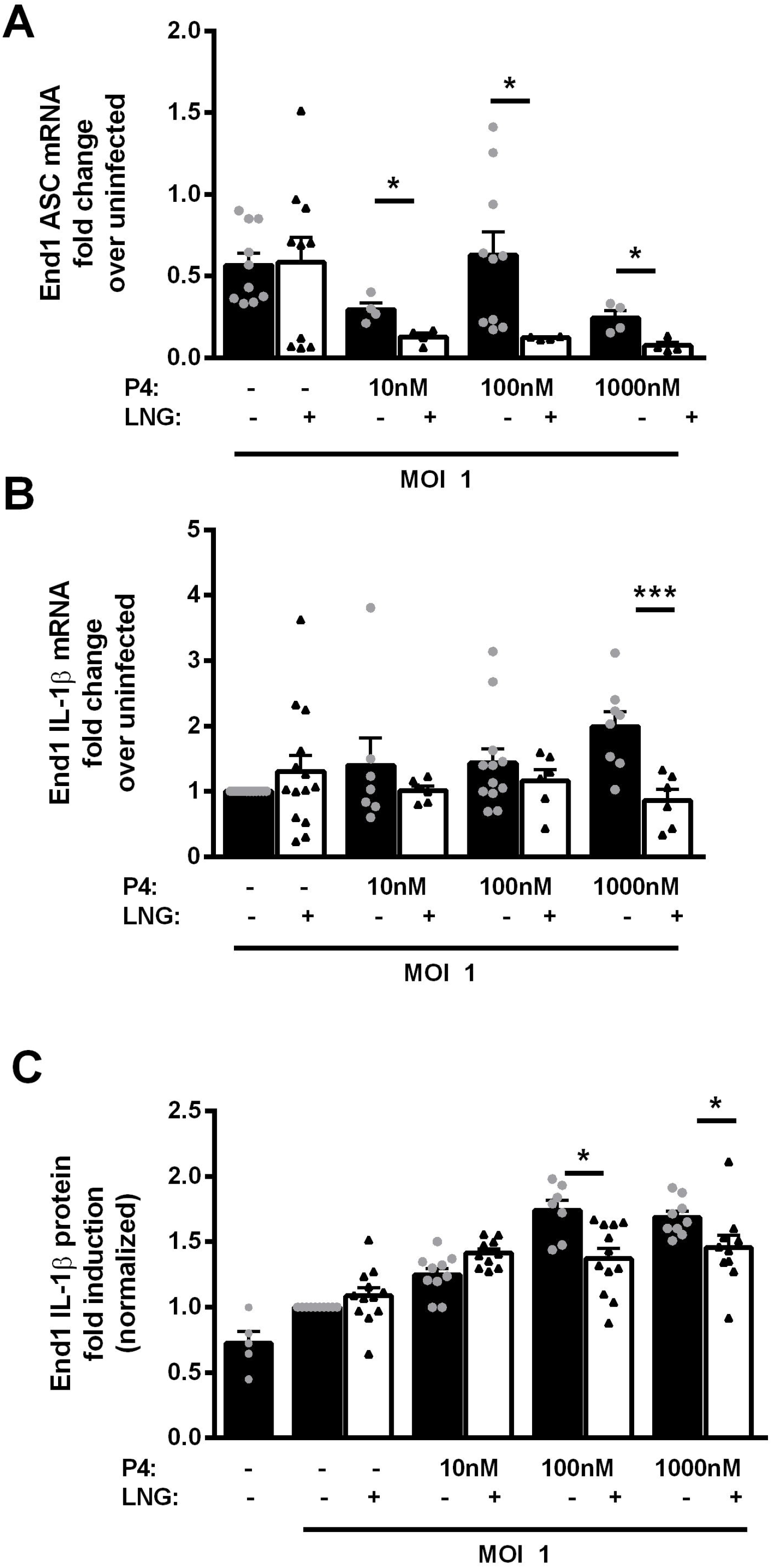
LNG suppresses endocervical epithelium inflammasome components and IL-1β during *C. trachomatis* infection. End1 cells were infected with Ct at an MOI of 1 and supernatants and RNA were harvested at 24 hours. (A) Asc mRNA levels were suppressed during Ct infection in all progesterone treatments, and this was significantly lowered by LNG addition. (B) IL-1β mRNA levels steadily increased during Ct infection in a dose-dependent manner with P4 addition, and this was suppressed by LNG addition at high levels of P4. (C) IL-1β protein level increased during Ct infection and this was also suppressed by LNG addition at high levels of P4. Data were analyzed by 2-way ANOVA. * p < 0.05, ** p < 0.01, *** p < 0.001.

We next moved from studying the immune response in structural cells to macrophages. In human THP.1 macrophage-like cells activated overnight with PMA at 25 nM, mRNA levels of ASC were upregulated in the absence of exogenous P4 but suppressed in the presence of LNG. Likewise at high concentrations of P4 treatment (100 and 1000 nM P4), P4 induced ASC mRNA while additional LNG treatment prevented this induction (Fig. 2A). Interleukin-1β mRNA was induced relative to baseline at all concentrations of P4, while addition of LNG again suppressed this induction (Fig. 2B). At the protein level, exogenous P4 induced IL-1β during Ct infection in a dose-dependent manner, and addition of LNG suppressed induction at high P4 concentrations (100 and 1000 nM) while trending towards suppression at lower concentrations (Fig. 2C).

**Figure 2.**
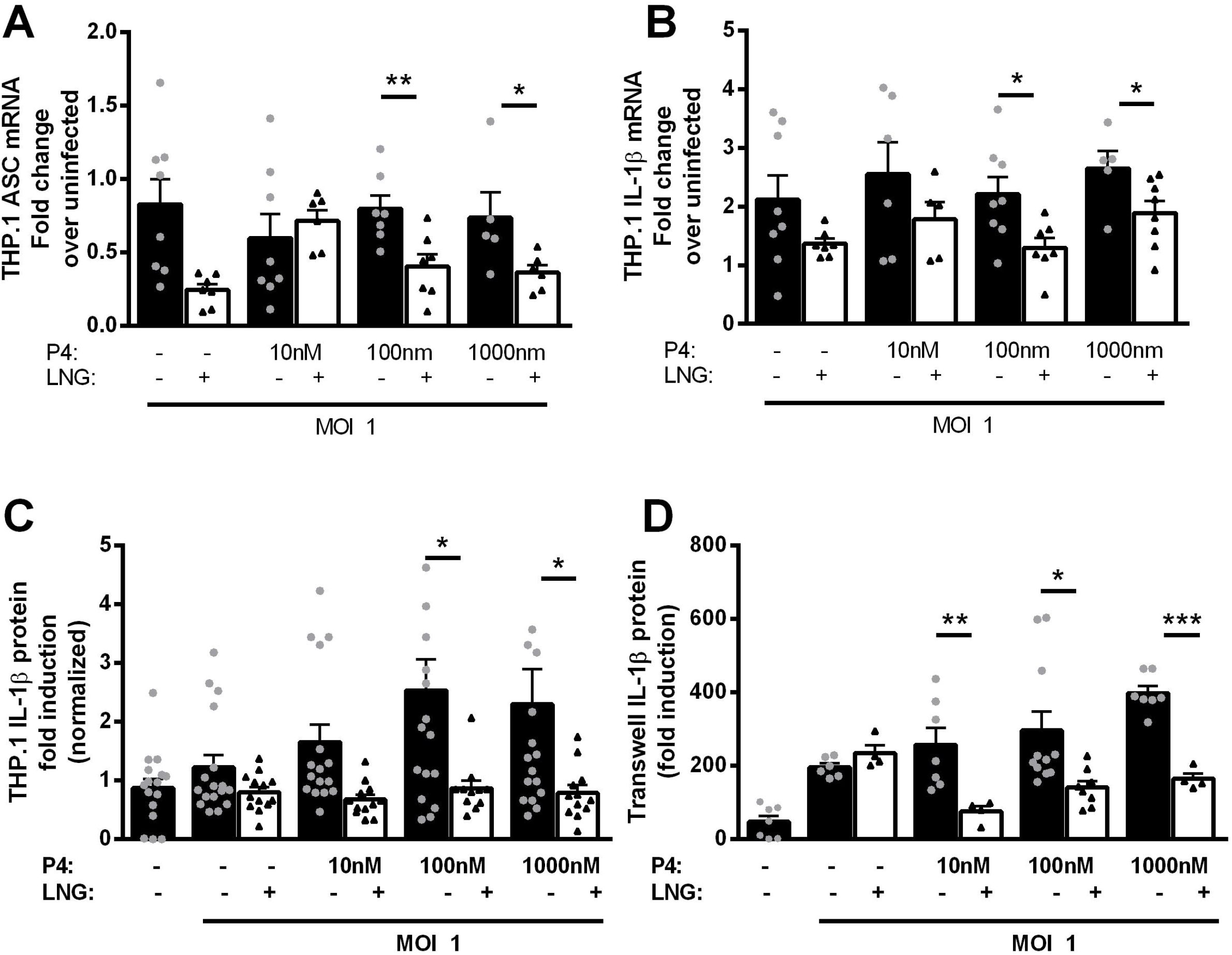
LNG suppresses macrophage inflammasome components and IL-1β during *C. trachomatis* infection and affects cellular communication between endocervical epithelium and immune cells. THP.1 cells were infected with Ct at an MOI of 1 and supernatants and RNA were harvested at 24 hours. (A) Asc mRNA levels were steady during Ct infection in all progesterone treatments, but this was significantly lowered by LNG addition. (B) IL-1β mRNA levels were also steady during Ct, but this was suppressed by LNG addition at high levels of P4. (C) IL-1β protein level increased during Ct infection and this was also suppressed by LNG addition at high levels of P4. (D) End1 cells were plated in the bottom chamber of a transwell setup and infected with Ct on day 1 with or without exogenous P4 and/or LNG and on day 2 cells were washed and THP.1 added to the top compartment of the transwell, and exogenous P4 and/or LNG added. LNG dramatically suppressed IL-1β protein from End1 and THP.1 co-cultures during Ct infection in all concentrations of exogenous P4. Data were analyzed by 2-way ANOVA. * p < 0.05, ** p < 0.01, *** p < 0.001.

In the body, endocervical cells are a primary target of Ct infection, and the ability of infected endocervical cells to activate or modulate a macrophage response may be important in disease progression. To model this, we plated End1 cells in the bottom of a transwell system, infected with Ct for 2 days in the presence or absence of increasing concentrations of exogenous P4 with or without LNG, then added THP.1 cells to the top chamber, separated by 2 μm porous membrane. In samples with only exogenous P4 added, IL-1β protein increased during Ct infection in a dose-dependent manner (Fig. 2D). Addition of LNG during infection prevented this increase in IL-1β protein in the presence of exogenous P4.

LNG can bind to numerous receptors, including the P4 receptor with high affinity, the glucocorticoid receptor with very low affinity, the mineralocorticoid receptor, and the androgen receptor. Employing pharmacological inhibitors of these various receptors, we lastly tested which receptor LNG could be acting through to suppress the IL-1β pathway during Ct infection. Using IL-1β protein as a readout, we treated End1 cells with RU-486 (P4 receptor inhibitor), CORT108297 (glucocorticoid receptor inhibitor), and ailanthone (androgen receptor inhibitor). Inhibition of the P4 receptor did not rescue LNG-mediated immune suppression during Ct infection (Fig. 3A). As expected, inhibition of the generally immunosuppressive glucocorticoid receptor raised all IL-1β responses to Ct infection, but also did not prevent the LNG-mediated immune suppression in the presence of P4 (Fig. 3B). However, addition of ailanthone prevented the LNG-mediated IL-1β suppression in both End1 cells (Fig. 3C) and in THP.1 cells (Fig. 3D), suggesting that LNG immunosuppression is mediated in part through the androgen receptor.

**Figure 3.**
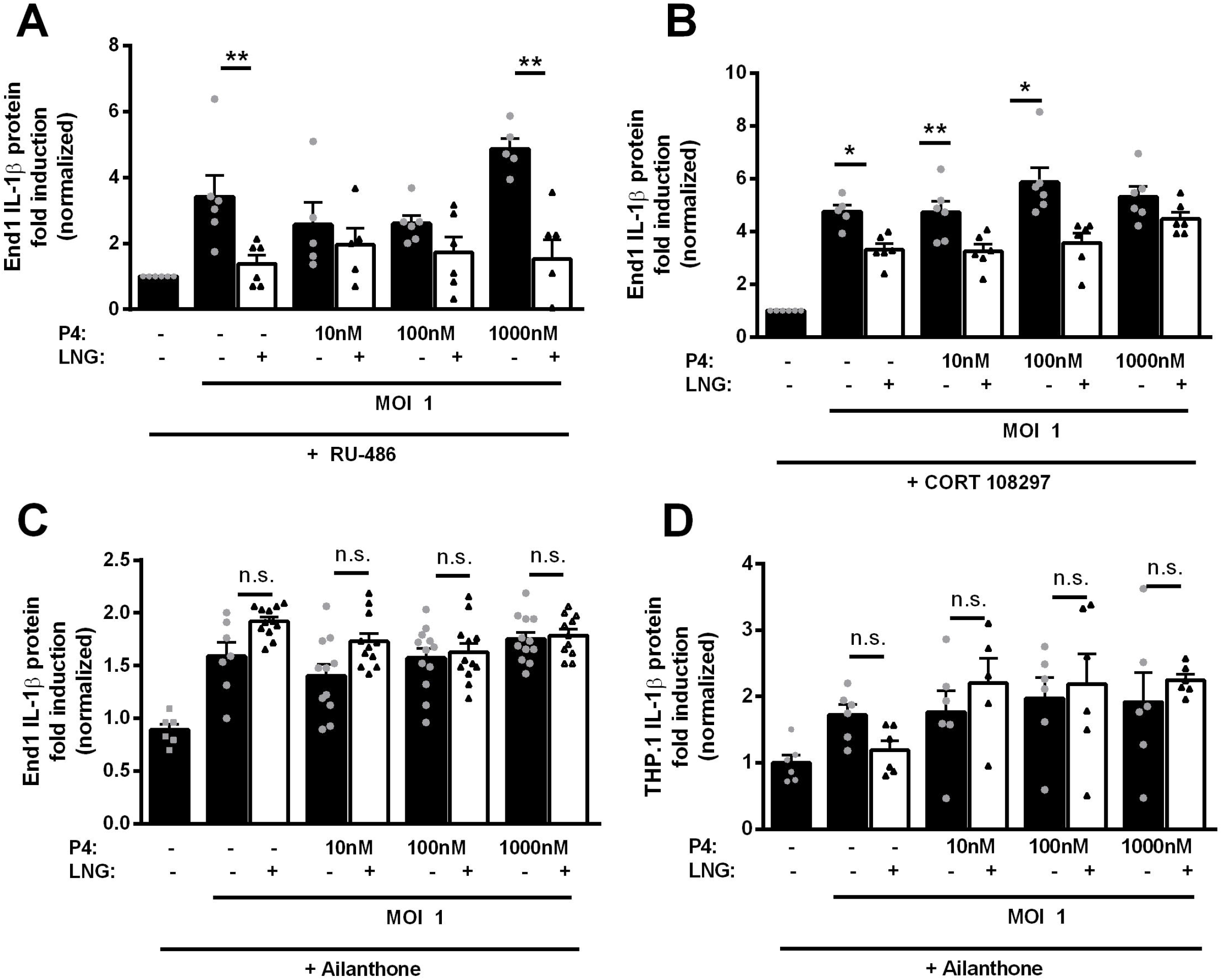
Immunosuppression by LNG appears to be tied to androgen receptor signaling. End1 cells (A-C) or THP.1 cells (D) were plated and infected with Ct in the presence of exogenous P4 and/or LNG and/or pharmacological inhibitors of known key LNG binding partners. Addition of RU-486, a progesterone receptor blocker (A) or CORT108297, a glucocorticoid receptor blocker (B), did not prevent LNG-mediated suppression of IL-1β responses during Ct infection. Addition of ailanthone, an androgen receptor blocker, rescued IL-1β production during Ct infection from both End1 cells (C) and THP.1 cells (D). Data were analyzed by 2-way ANOVA. * p < 0.05, ** p < 0.01, *** p < 0.001.

To address the generalizability of our findings, we repeated selected experiments using the TLR2 agonist PAM3CSK4 instead of Ct. TLR2 stimulation of THP.1 macrophages by PAM3CSK4 at 1 uM resulted in modest induction of IL-1β. LNG treatment in the absence of progesterone did not change this induction. However, addition of P4 in increasing levels resulted in upregulation of IL-1β while co-treatment of these cells with LNG suppressed P4-mediated upregulation of IL-1β.

In this study we have shown that the effects of LNG are not always apparent in the absence of other biological variables such as hormone. In the absence of exogenous P4, LNG often failed to suppress IL-1β in response to Ct infection in both endocervical cells and macrophages. However, in the presence of P4, LNG suppressed the IL-1β response from both these cell lines individually and, more dramatically, when co-cultured in a transwell system.

The limitations of this study merit discussion. The effect size in certain treatments was small, as little as 1.5 fold induction over baseline. This has been observed before^33^, and consistent with their findings, treatment with a TLR agonist induced a greater IL-1β response than Ct alone (Fig. S1), which was nonetheless suppressed by co-treatment with LNG. Additionally, the concentration of LNG to use was extrapolated from tissue to cell culture medium based on early LNG-IUS models, which released 30 ng instead of 20 ng of LNG per day as they do now; to our knowledge, there are no more recent studies determining the concentration of LNG in the uterus during modern LNG-IUS use. The extrapolation is also imperfect, based on pg/mg protein, then scaled to the amount of protein obtained from a well of endocervical cells. However, 100 ng/ml is likely low enough to be within the confines of a biologically relevant concentration.

We found that in the absence of exogenous P4, LNG often had little effect on IL-1β levels, and exogenous P4 was necessary to observe immunosuppression by LNG: it is unclear why the presence of progesterone is necessary for LNG-mediated immune suppression via the androgen receptor. It is possible that P4 levels induce androgen receptor expression, or that in this system exogenous P4 is sufficiently concentrated to compete with LNG for P4 receptor binding sites, thus pushing LNG to bind androgen receptor as the next preferred receptor.

Finally, there is the question of whether this LNG-mediated suppression of IL-1β is beneficial to the host or detrimental. Abdul-Sater, *et al.* showed in 2009 that intracellular growth of Ct is stimulated by caspase-1 activation^16^, and IL-1β has been shown to contribute to fallopian tube scarring^15^. Inhibition of IL-1β may limit Ct spread and damage. However, other research has shown a critical role for IL-1β in Ct infection^33^.

In summary, the interaction of exogenous LNG in the uterus combined with endogenous P4 cycling merits further investigation into the interaction of these progestins, especially during innate immune responses to pathogens. The mechanism of androgen receptor-mediated IL-1β immunosuppression likewise requires further study. These data suggest that LNG-mediated immunomodulation could impact women seeking or using an LNG-based hormonal contraceptive who are in high-risk groups for contracting STIs, and could result in recommendations for more frequent STI screening.

## Materials and methods

### Reagents

A clinical isolate of *C. trachomatis* serovar E was purchased from the University of Washington (Seattle, WA, USA). The End1 endocervical cell line and THP.1 monocyte cell line were purchased from ATCC (Manassas, Virginia, USA). End1 was grown in Keratinocyte serum-free media purchased from Gibco/Thermo Fisher (Waltham, Massachusetts, USA), and THP.1 cells were grown in RPMI-1640 supplemented with 10% heat-inactivated fetal bovine serum without antibiotics (Gibco). P4, LNG, RU-486, CORT108297, ailanthone, and PAM3CSK4 were purchased from Sigma-Aldrich (St. Louis, MO, USA). Reverse transcription was performed using the Qiagen QuantiTect RT kit (Germantown, MD, USA), and qPCR was performed using the Radiant SYBR Green Lo-ROX qPCR kit (Alkali Scientific, Inc, Fort Lauderdale, FL, USA). Sequences for probes were as follows: 18s FWD TGGAGGATGGACCACATCAG, REV CTGGAGCACTTTGCGTCGTT; ASC FWD TGGATGCTCTGTACGGGAAG, REV CCAGGCTGGTGTGAAACTGAA; IL-1β FWD GCCAGGAACCATGCTTTGAC, REV CTCGGGATTCTTTCAAGCCCT. ELISAs were performed using the R&D human DuoSet IL-1β ELISA kit (Minneapolis, MN, USA).

### Cell culture

End1 cells were cultured in 96-well plates or 24-well dishes at 1 × 10^5^ cells/well or 2.5 × 10^5^ cells/well, respectively. P4 was added prior to Ct infection at 10, 100, and 1000 nM, as was LNG at a concentration of 100 nM. Ct was added at an MOI of 1, infected for 48 hrs, and supernatants harvested for protein analysis by ELISA and monolayers harvested for mRNA analysis by qPCR according to manufacturers’ instructions.

### Statistical analysis

Experiments were repeated 3 times to ensure reproducibility. Each experiment had 2-6 replicates per condition. qPCR reactions were run in duplicate. Data was analyzed by 2-way ANOVA using GraphPad Prism version 6 (La Jolla, CA, USA).

---

**Figure S1. LNG suppresses P4-mediated increase in IL-1β during stimulation with TLR2 agonist PAM3CSK4. THP.1 cells were stimulated with increasing amounts of P4 with or without LNG, and exposed to the TLR2 agonist PAM3CSK4**. IL-1β levels increased during TLR2 stimulation in the presence of increasing P4, but this increase was blocked by LNG addition. * p < 0.05.

