## Supplementary figures and images for "Levonorgestrel modulates innate immune activation in human endocervical epithelium and monocytes during *Chlamydia trachomatis* infection"

### Figure S1

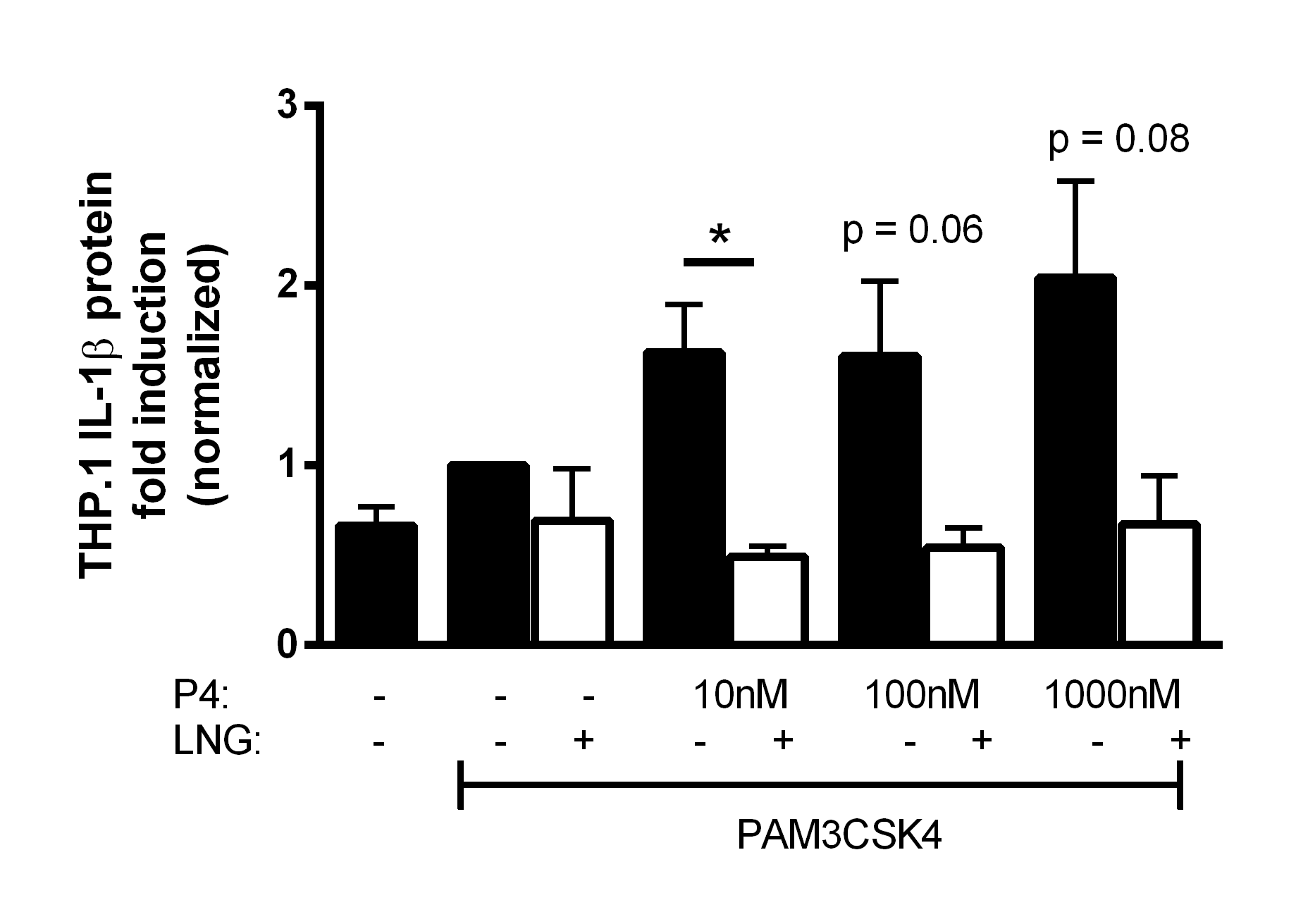
